# Divergent *In vitro* MIC Characteristics and underlying isogenic mutations in host-specialized *Pseudomonas aeruginosa*

**DOI:** 10.1101/118109

**Authors:** Xuan Qin, Alexander L Greninger, Chuan Zhou, Amanda Adler, Shuhua Yuan, Danielle M. Zerr

## Abstract

Clinical isolates of *Pseudomonas aeruginosa (Pa)* from patients with cystic fibrosis (CF) are known to differ from those associated with infections of non-CF hosts in colony morphology, drug susceptibility patterns, and genomic hypermutability. Although *Pa* isolates from CF have long been recognized for their overall higher resistance rate calculated generally by reduced “percent susceptible”, this study takes the approach to compare and contrast Etest MIC distributions between two distinct cohorts of clinical strains (n=224 from 56 CF patients and n=130 from 68 non-CF patients respectively) isolated in 2013. Logarithmic transformed MIC (logMIC) values of 11 antimicrobial agents were compared between the two groups. CF isolates tended to produce heterogeneous and widely dispersed MICs compared to non-CF isolates. By applying a test for equality of variances, we were able to confirm that the MICs generated from CF isolates against 9 out of the 11 agents were significantly more dispersed than those from non-CF (p<0.02-<0.001). Quantile-quantiles plots indicated little agreement between the two cohorts of isolates. Based on whole genome sequencing of 19 representative CF *Pa* isolates, divergent gain- or loss-of-function mutations in efflux and porin genes and their regulators between isogenic or intra-clonal associates were evident. Not one, not a few, but the net effect all adaptive mutational changes in the genomes of CF *Pa*, both shared and unshared between isogenic strains, are responsible for the divergent heteroresistance patterns. Moreover, the isogenic variations are suggestive of a bacterial syntrophic lifestyle when “lockedȍ inside a host focal airway environment over prolonged periods.

**Significance statement:** Bacterial heteroresistance is associated with niche specialized organisms interacting with host species for prolonged period of time, medically characterized by “chronic focal infections”. A prime example is found in *Pseudomonas aeruginosa* isogenic/non-homogeneous isolates from patient airways with cystic fibrosis. The development of pseudomonal polarizing MICs *in vitro* to many actively used antimicrobial agents among isogenic isolates and “Eagle-type” heteroresistance patterns are common and characteristic. Widespread isogenic gene lesions were evident for defects in drug transporters, DNA mismatch repair, and many other structural or cellular functions—a result of pseudomonal symbiotic response to host selection. Co-isolation of extremely susceptible and resistant isogenic *Pa* strains suggests intra-airway evolution of a multicellular syntrophic bacterial lifestyle, which has laboratory interpretation and clinical treatment implications.

## Introduction

In the airway environment of patients with cystic fibrosis (CF), isolates of *Pseudomonas aeruginosa (Pa)* and other common bacterial cohabitants undergo host airway niche adaption and virulence attenuation (1, 2). As soon as the organism enters the CF airway, it is under enormous host selective pressure that is reflected by pseudomonal transition into a hypermutable state with loss of function mutations in *mutS* and *mutL* (3–6). The diverse MIC patterns generated among co-isolates are in part a result of pseudomonal isogenic divergence in response to host and antimicrobial pressure (3, 7). Similarly, bacterial small colony variants (SCV) have been described to coexist as isogenic variants *in vitro* (8). Bacterial SCVs found in clinical isolates of *Staphylococcus aureus, Enterococcus faecium, Stenotrophomonas maitophiiia, Escherichia coli*, and *Pa* are characteristically associated with chronic infections of the bone/joint, urinary tract, sinus/ear, etc., and do not as predictably respond to antibiotic treatment (8–14). Infections involving isogenic divergent organisms are frequently prolonged and localized without bloodstream dissemination, and are unable to live outside of the chronically affected host focal environment and therefore have not been shown to be able to transmit between patients (15–18). With advances in treatment of malignances and other complex diseases, there is a growing patient population receiving frequent and long-term antibiotic therapy that may be at increased risk of developing infections involving isogenic divergent organisms. The so-called “persister” organisms frequently show “heteroresistance” that describes a phenomenon where subpopulations of isogenic bacteria exhibit a range of susceptibilities to a particular antibiotic agent (19). Clinical laboratories are currently not equipped or adequately guided for appropriate testing or interpretation of non-homogeneous bacterial isolates or co-isolates of clonal associates that produce highly variable and isogenic heteroresistance patterns (3, 19, 20).

Antibiograms are routinely constructed by clinical microbiology laboratories on an annual basis (21). The format of hospital antibiograms is standardized by reporting “percent susceptible” using interpretive criteria recommended by professional advisory organizations such as Clinical Laboratory Standard Institute (CLSI) (22). Additionally, the rules used for construction of institutional antibiograms require the inclusion of “first isolate during the time period analyzed”, which refers to the initial microbial isolate of a particular species recovered from a patient, regardless of culture source or antimicrobial profile (22). This rule clearly has limitations in the setting of isogenic CF *Pa* stains, given the phenotypic diversity and ongoing evolution of resistance profiles (7, 23). Unlike the pseudomonal isolates from non-CF patients, *Pa* isolates from repeat cultures from CF patients usually do not have consistent phenotypes (based on colony morphology or susceptibilities) in repeat cultures even a few days apart. Isogenic strains tend to produce visually diverse colony variants accompanied by respective divergent or heterogeneous resistance patterns (7,19). Moreover, common laboratory practice emphasizes the isolation and purification of a phenotypically homogenous bacterial stain(s) before susceptibility testing, which has limited our ability to understand bacterial organisms that are no longer assuming single cell replicative mode of living, but rather an isogenic interdependent multicellular mode of replication (7).

In our previous study, we found that co-isolation of patient-specific isogenic *Pa* strains with widely divergent MICs to sulfamethoxazole-trimethoprim (SMX-TMP) was common in CF airways specimens (7). Moreover, isogenic lesions in the responsible efflux apparatus *mexAB-oprM* were found in clonally coexisting *Pa* isolates. Once again, the most visited resistance-nodulation-cell division (RND) family of multi-drug efflux apparatus MexAB-OprM, a prominent drug-proton antiporter and its immediate transcription regulator MexR, is responsible for such intra-clonal divergent SMX-TMP MICs (24–26). Our study also showed the isogenic isolates were often auxotrophic for either arginine and/or methionine. We have also demonstrated that nutritional cross-feeding supported co-growth between prototroph and auxotroph complements with intra-clonal preferences; thus we postulate that multicellular resistant syntrophy is the mechanism of pseudomonal persistence without clearance and serendipitously, the absence of repeat attacks thereafter by clonally unrelated wild-type *Pa* strain(s) in CF airways (3, 27). High level resistance to aztreonam and carbapenems in CF *Pa* has long been be linked to inactivation or alteration of a drug transporter porin D and many adaptive mutations in regulatory determinants, such as *mexR, nalC* and *nalD*, contributing to *mexAB-oprM* hyper-expression (28–30).

This investigation aimed to describe heteroresistance patters in CF *Pa* isolates and, compare and contrast the characteristic antimicrobial MIC distributions between two cohorts of *Pa* isolates (non-CF and CF) in order to confirm pseudomonal divergent susceptibilities *in vitro* and syntrophic resistance (or heteroresistance) mechanisms inferred from distinct *in vitro* MIC patterns and diverse mutations in corresponding gene determinants.

## Results

### Heteroresistance generated from seemingly homogeneous isolates *in vitro*

To demonstrate heteroresistance patterns of CF *Pa* isolates at the seemingly “homogeneous” strain level, we have summarized 4 examples of this phenomenon in Figure 1. The first example can be common to both CF and non-CF *Pa* isolates where “inner colonies” were growing above the meropenem MIC “cut-point” of the predominant growth from a seemingly pure clone of an isolate (Figure 1a). A meropenem MIC result of 32μg/μl was reported for this isolate. The second example is more common in CF *Pa* isolates (Figure 1b). In this example, a CF *Pa* isolate showed clear heteroresistance against ceftazidime, in which two subpopulations of the same strain are generating two distinct MICs at either 0.5μg/μl or 32μg/μl, depending on whether the “cut-point” is accepted at “80%” or “100%” inhibition of the bacterial growth, respectively. A ceftazidime MIC of 32μg/μl was reported for this isolate. The third example of heteroresistance is also common in CF *Pa* (Figure 1c). This example shows enhanced growth within a range of amikacin concentrations and the most prominent bacterial growth was at the amikacin concentration of 4-96μg/μl. An amikacin MIC of 96μg/μl was reported for this isolate. The final example has been only found in CF *Pa* where the bacterial growth was most prominent at the higher ticarcillin-clavulanate range of >2μg/μl (Figure 1d). A ticarcillin-clavulanate MIC of >256 μg/μl was reported for this isolate. Although they occur less frequently, the third and forth types of heteroresistance would not have been recognized without a manual broad range broth dilution series, or Etest agar diffusion methods *in vitro.* Moreover, all four types of heteroresistance patterns have been highly reproducible when the areas of the growths producing heterogeneous growth properties to the corresponding antibiotic agent(s) were further subcultured and “purified” to repeat against the same agent (data not shown).

**Figure 1.**
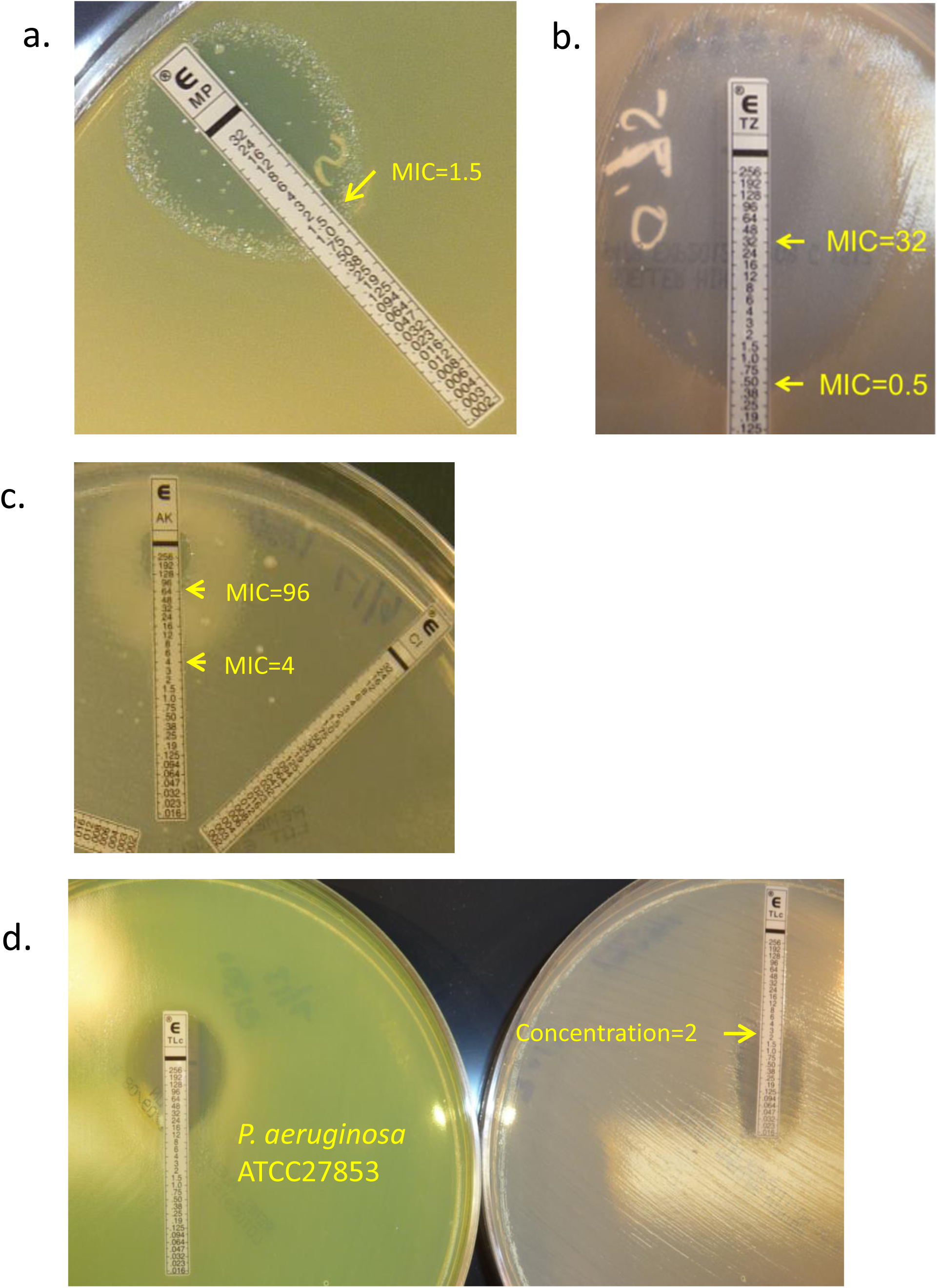
*P. aeruginosa* “strain-specific” heteroresistance. (A) inner colonies growing above the meropenem MIC cut point, (B) two subpopulations of the same strain generating two distinct ceftazidime MICs, (C) enhanced bacterial growth within a range of amikacin concentrations, (D) bacterial growth that is most prominent at the higher ticarcillin-clavulanate range. MIC concentration μg/μl.

### Bacterial isolates from cultures included for antimicrobial MIC analysis

A total of 354 clinical *Pa* strains were included in this analysis: 224 CF *Pa* isolates from 100 cultures from 56 patients and 130 non-CF *Pa* isolates from 99 cultures from 68 patients (Table 1). The CF *Pa* isolates were generally from sputum samples, oropharyngeal swabs, and bronchoalveolar lavage specimens. The non-CF *Pa* strains came from variety of specimens types (tracheal aspirates n=59, bronchoalveolar lavage n=5, sputum n=1, urine n=28, wound n=6, external ear n=9, blood n=5, cerebral spinal fluid n=3, catheter tip n=1 sinus n=2, stool (hematology-oncology patients) n=7, other n=4.

**Table 1.** Number of *P. aeruginosa* isolates associated with the number of cultures and patients.

|  | CF<br>*P. aeruginosa* | Non-CF<br>*P. aeruginosa* | P value |
| --- | --- | --- | --- |
| Number of patients | 56 | 68 | -- |
| Number of cultures | 100 | 99 | -- |
| Number of isolates | 224 | 130 | -- |
| Number (%) of cultures with >1 isolates | 64 (64%) | 28 (28%) | -- |
| Number of cultures with >1 isolate with discordant susceptibilities<sup>b</sup> | 41 (64%) | 6 (21%) | -- |
| Average isolates per culture<sup>a</sup> | 2.24±1.19 (1-5) | 1.31±0.51 (1-3) | <0.001 |
| Average number of agents involved in discordant susceptibilities per culture<sup>b</sup> | 1.73±2.14 (0-10) | 0.39±0.83 (0-3) | 0.001 |

-- Not evaluated. <sup>a</sup> Results reported as mean and standard deviation (range). <sup>b</sup>Discordant susceptibilities include only those with changes in interpretations between susceptible and resistant categories. Changes with MICs producing between intermediate and susceptible or resistant are not included.

Overall, 64% of cultures from CF airway samples had ≥2 *Pa* colony types compared to only 28% of non-CF specimens (Table 1). The *Pa* co-isolates from the CF cohort cultures were more likely to have discordant susceptibilities to at least one antimicrobial agent (64%) compared to coisolates from non-CF cultures (21%). Similarly, the average number of *Pa* colony types per culture are significantly higher in CF cohort versus the non-CF cohort (2.24 versus 1.31; p<0.001), and the average number of agents associated with discordant susceptibilities was also higher in the CF cohort (1.73 vs 0.39; p=0.001, Table 1).

### Antimicrobial susceptibility by percent susceptible

Based on CLSI MIC breakpoint criteria, antimicrobial susceptibilities of the *Pa* isolates against the 11 agents tested by Etest MIC method are compared and summarized in Table 2. The proportion of *Pa* isolates susceptible to any of the 3 aminoglycoside agents – amikacin, gentamicin, and tobramycin – is significantly lower in CF *Pa* than those of non-CF *Pa* (p <0.001 for all three comparisons). Similarly, the proportion of *Pa* isolates susceptible to cefepime and ciprofloxacin was also significantly lower in CF *Pa* (p <0.001 and p=0.002, respectively). In contrast, the proportion of *Pa* isolates susceptible to sulfamethoxazole-trimethoprim is significantly higher in CF *Pa* than that of non-CF *Pa* (p <0.001). The proportion of isolates susceptible to other agents did not differ between the two cohorts (Table 2).

**Table 2.** Antimicrobial susceptibilities presented by percent susceptible.

|  | CLSI MIC breakpoints |  |  | Etest range<br>(µg/µl) | CF *Pa*<br>Percent<br>susceptible<br>(n=224) | non-CF *Pa*<br>Percent<br>susceptible<br>(n=130) | p<br>value |
| --- | --- | --- | --- | --- | --- | --- | --- |
|  | S | I | R |  |  |  |  |
| Amikacin | ≤16 | 32 | ≥64 | 0.016 - 256 | 68 (30%) | 117 (90%) | <0.001 |
| Gentamicin | ≤4 | 8 | ≥16 | 0.064 - 1024 | 49 (22%) | 97 (75%) | <0.001 |
| Tobramycin | ≤4 | 8 | ≥16 | 0.064 - 1024 | 144 (64%) | 123 (95%) | <0.001 |
| Aztreonam | ≤8 | 16 | ≥32 | 0.016 - 256 | 186 (83%) | 105 (82%) | 0.81 |
| Ceftazidime | ≤8 | 16 | ≥32 | 0.016 - 256 | 190 (85%) | 111 (85%) | 0.89 |
| Cefepime | ≤8 | 16 | ≥32 | 0.016 - 256 | 108 (48%) | 108 (83%) | <0.001 |
| Imipenem | ≤2 | 4 | ≥8 | 0.002 - 32 | 97 (43%) | 53 (41%) | 0.64 |
| Meropenem | ≤2 | 4 | ≥8 | 0.002 - 32 | 178 (79%) | 108 (83%) | 0.41 |
| PTc | ≤16/4 | 32/4-64/4 | ≥128/4 | 0.016 - 256 | 185 (83%) | 111 (85%) | 0.49 |
| Ciprofloxacin | ≤1 | 2 | ≥4 | 0.016 - 32 | 169 (75%) | 116 (89%) | 0.002 |
| SMX-TMP | ≤2/38 |  | ≥8/152 | 0.002 - 32 | 82 (37%) | 24 (18%) | <0.001 |

CF, cystic fibrosis; *Pa*, *Pseudomonas aeruginosa*; I, intermediate; PTc, piperacillin-tazobactam; R, resistant; S, susceptible; SMX-TMP, sulfamethoxazole-trimethoprim.

### Heterogeneous susceptibilities associated with CF *Pa* using histograms and quantile-quantile plots generated from logMIC

When compared to logMIC distribution patterns generated from non-CF *Pa*, antimicrobial logMICs generated from the CF *Pa* isolates are generally more widely dispersed over the Etest MIC ranges for most of 11 agents tested (Fig 2). Based on quantile-quantile plots of the logMICs from the two cohorts, antimicrobial MICs generated by CF *Pa* are likely to produce more extreme values at both the higher and lower ends, the so-called “heavy tails” of the measurable range (Fig 2). Moreover, there is little agreement between the distributions of logMICs generated from isolates between CF and non-CF cohorts when analyzed by the quantile-quantiles plots (Fig 2).

**Figure 2.**
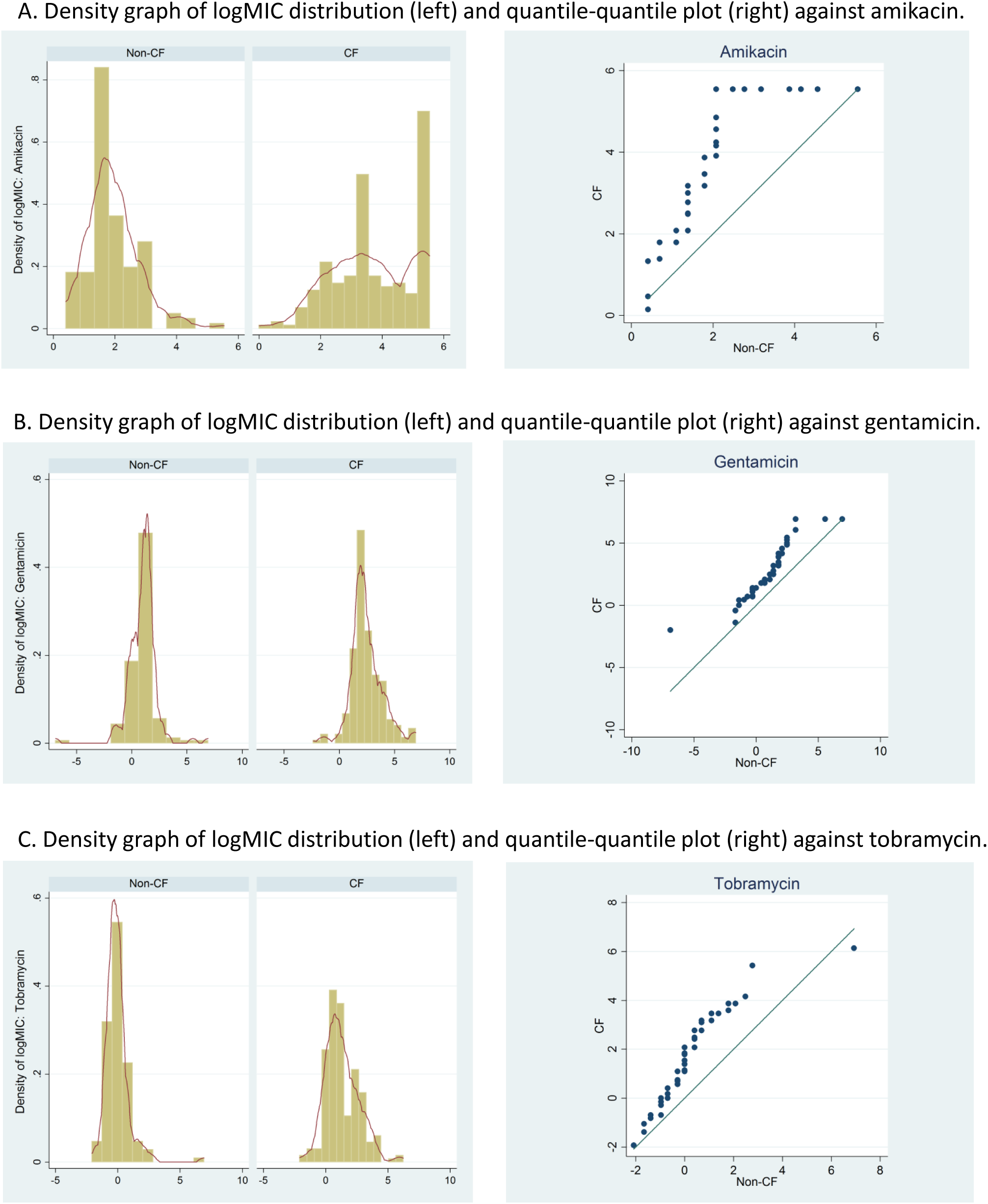
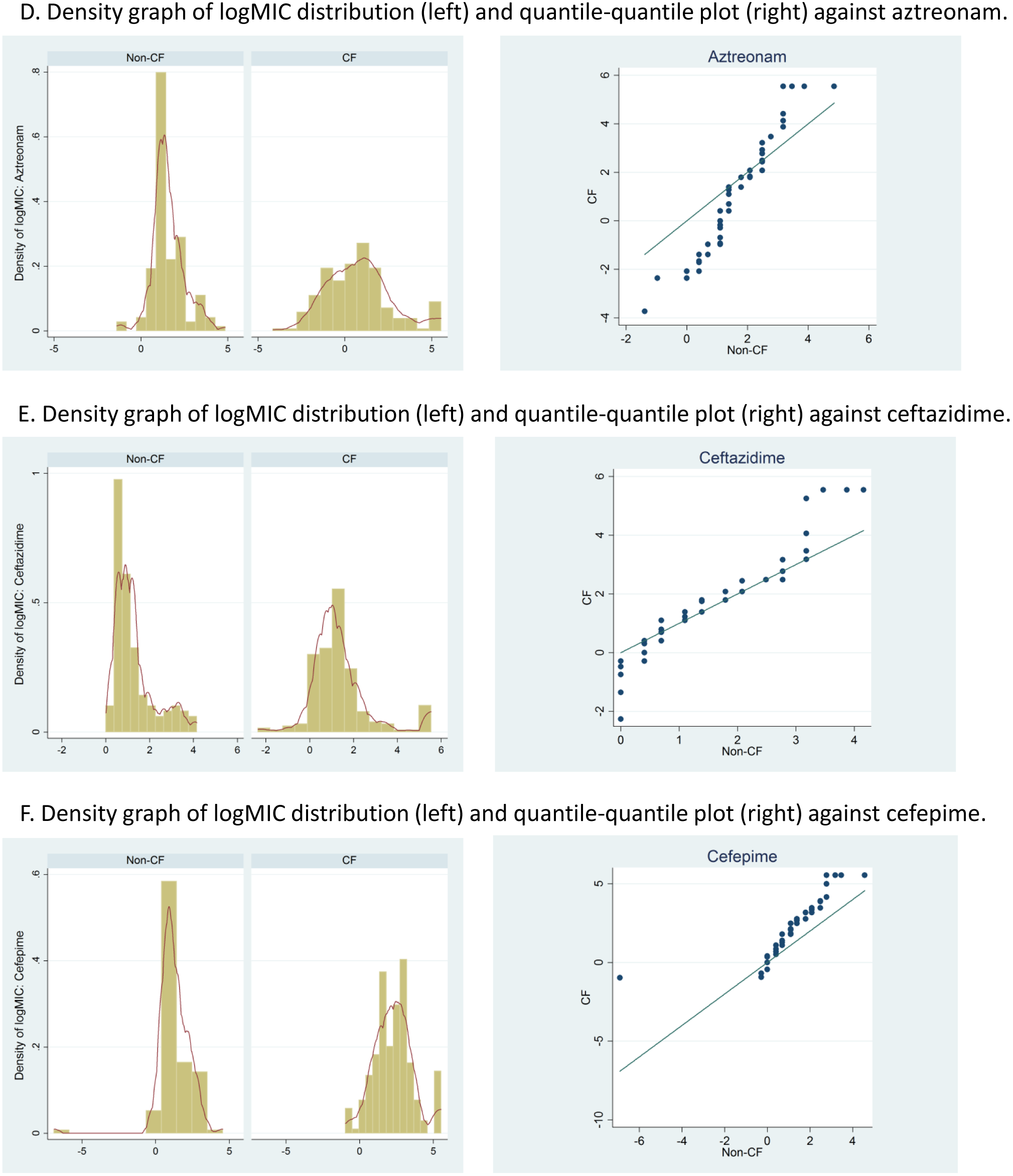
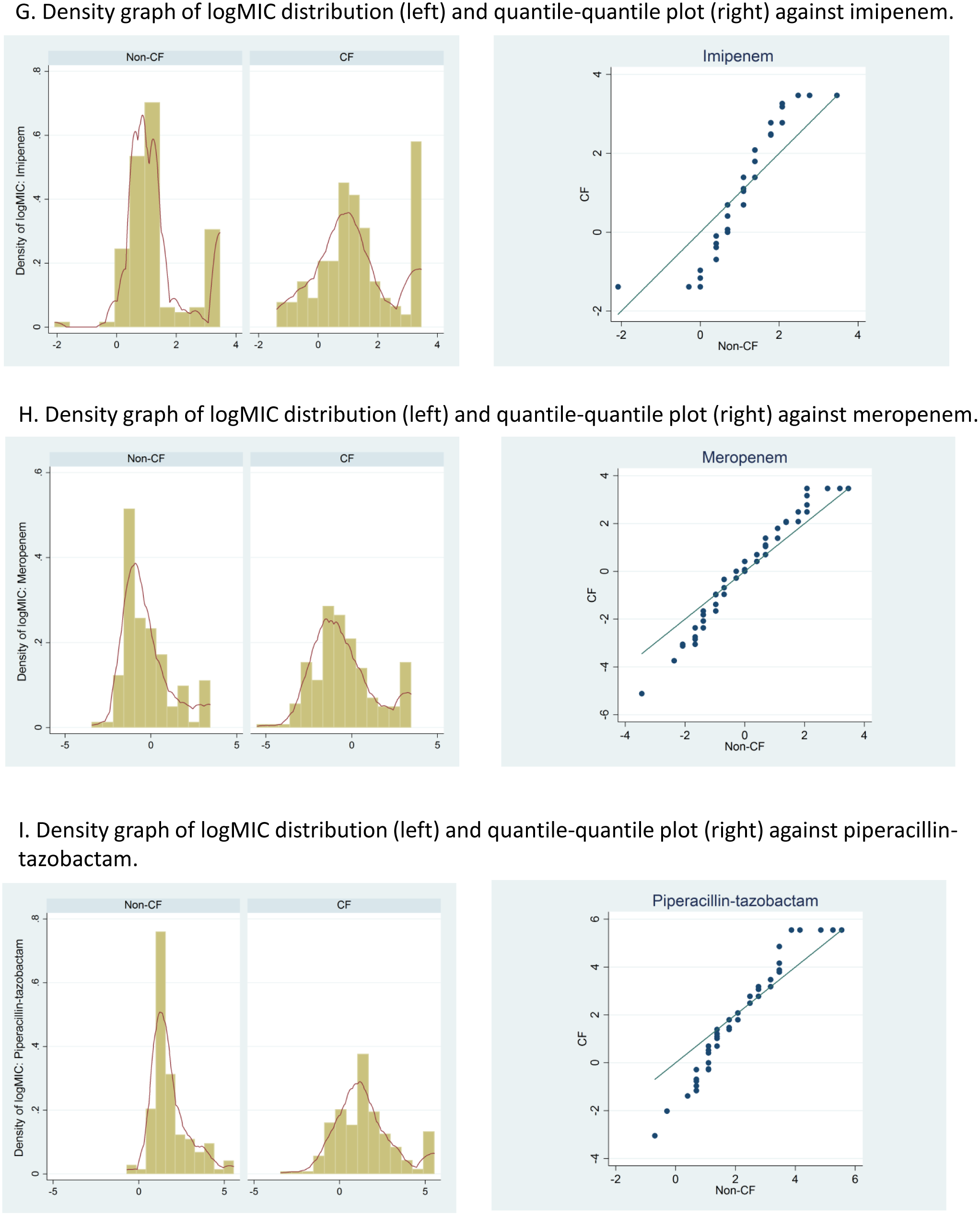
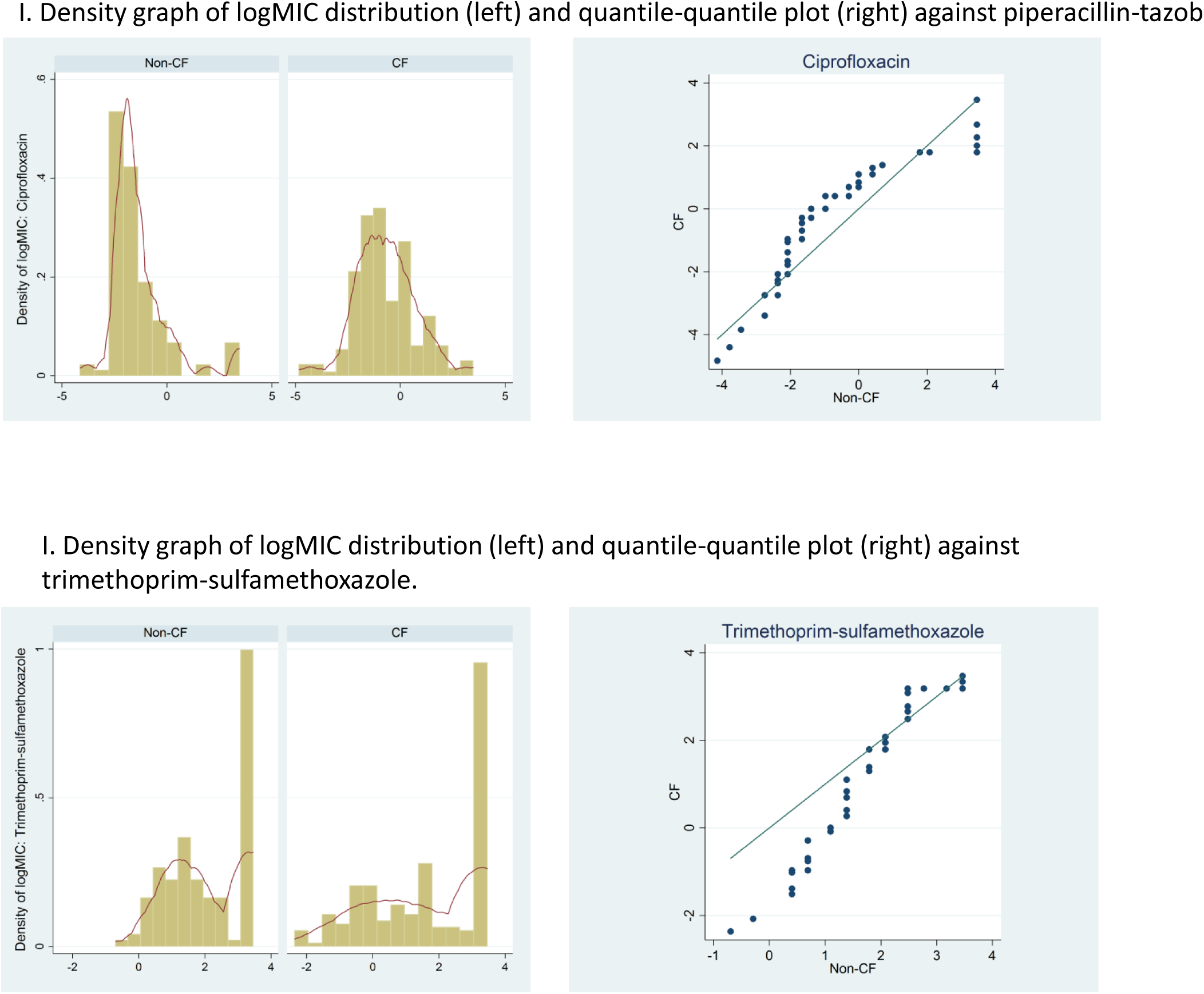
Tobramycin and ceftazidime logMIC distributions of *P. aerugionsa* isolates from patients with and without cystic fibrosis. The left side of the figure demonstrates the density graphs of logMIC distribution and quantile-quantile plots are on the right. Antimicrobial logMIC distributions of *P. aerugionsa* isolates from patients with and without cystic fibrosis

The test for equality of variances demonstrated that CF pseudomonal MIC distribution were highly skewed compared to the non-CF (Fig 2). The logMIC distributions for nine out of 11 antimicrobials tested on CF isolates were significantly more dispersed compared to the logMIC distributions generated from non-CF isolates (p = <0.02 − <0.001; Table 3).

**Table 3.** Test for equality of variances between MICs in cystic fibrosis or non-cystic fibrosis *Pseudomonas aeruginosa* isolates.

| Antimicrobial | Standard Deviation of Non-CF | Standard Deviation of CF | p-value |
| --- | --- | --- | --- |
| Amikacin | 0.85 | 1.4 | <.001 |
| Gentamicin | 1.33 | 1.45 | 0.06 |
| Tobramycin | 0.99 | 1.33 | <.001 |
| Aztreonam | 0.92 | 1.92 | <.001 |
| Ceftazidime | 0.93 | 1.39 | 0.02 |
| Cefepime | 1.13 | 1.37 | <.001 |
| Imipenem | 1.07 | 1.37 | <.001 |
| Meropenem | 1.45 | 1.88 | 0.002 |
| PTc | 1.12 | 1.78 | <.001 |
| Ciprofloxacin | 1.42 | 1.43 | 0.09 |
| SMX-TMP | 1.19 | 1.75 | <.001 |

CF, cystic fibrosis; PTc, piperacillin-tazobactam; SMX-TMP, sulfamethoxazole-trimethoprim

### Whole genome sequencing of isogenic CF *Pa* strains showing polarizing MICs *in vitro*

A total of 19 CF *Pa* isolates belonging to 9 patients were selected for whole genome sequencing. All patient co-isolates selected for genome sequencing were either from the same cultures (patient 4, 6, 9,19, 34, 51 and 53) or from different cultures of the same host (patient 5 and 28) (Table 4). When sequence determinants associated with CF *Pa* strains were compared, the polymorphisms found in common to those of PA14 were treated as “neutral” changes for study strains assuming those may not result in major MIC discrepancies or other CF *Pa* phenotypic characteristics in reference to those generated from PAO1 (Table S1-S4).

**Table 4.**
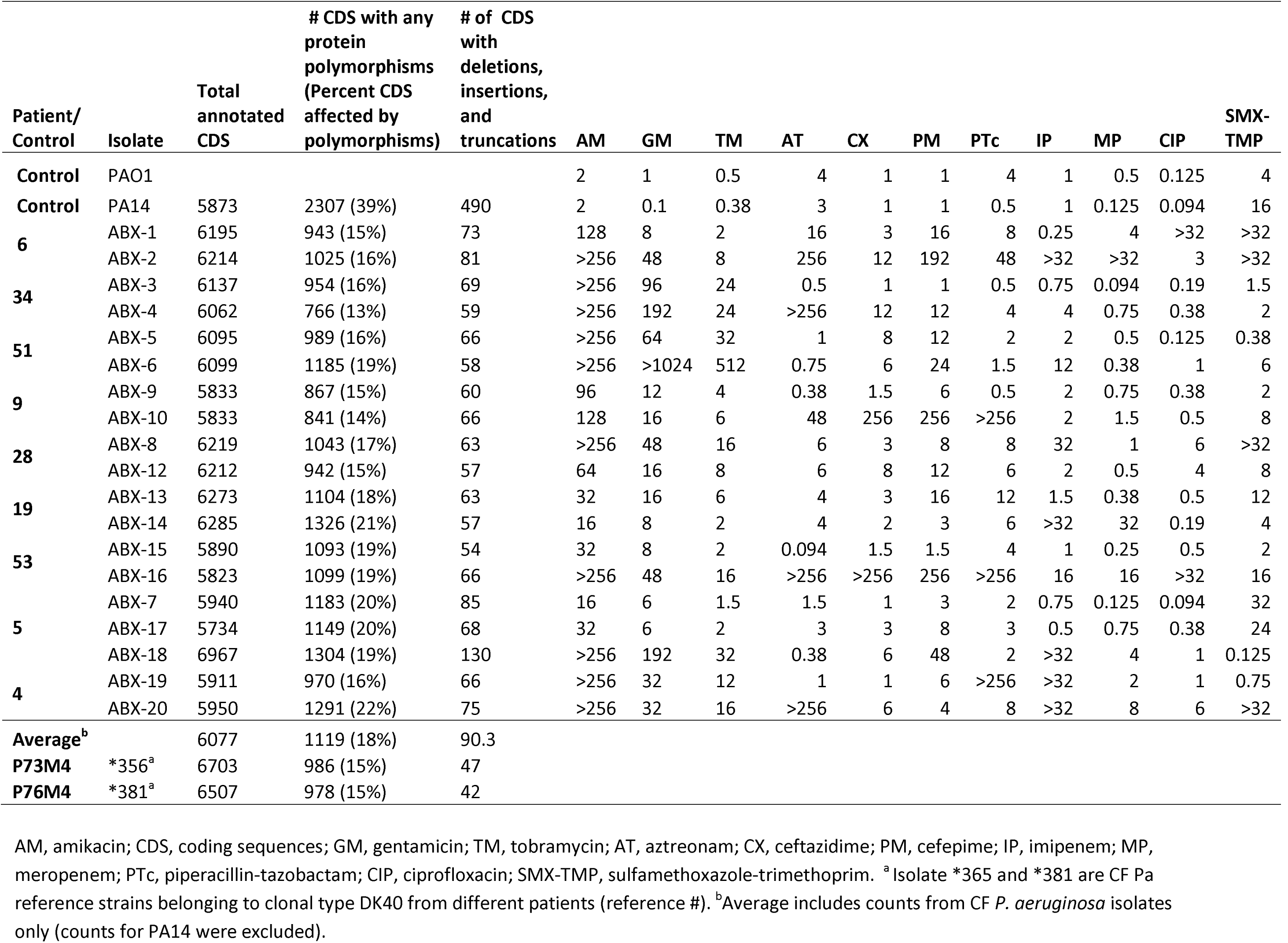
Summary of *in vitro* antimicrobial MICs and overall genomic polymorphisms of the 19 CF *P. aeruginosa* isolates and control strains.

The total number of DNA coding sequences (CDSs) affected by polymorphic mutations relative to the PAO1 reference (average 6088, range from 5734 to 6967, including insertions, deletions, truncations, and amino acid substitutions) or percent of CDSs affected among the 19 CF *Pa* isolates did not vary significantly – an average of 18% (range from 13% to 21%, Table 4). While the reference *Pa* genomes ^*^356 and ^*^381 were more closely related to PA14, the ABX isolates were showing closer distance to PAO1 according to pan genome tree (Fig 3). The isogenic relationship between co-isolates appeared to be patient-specific and clonally related (Fig 5 and Table S1-S4, a similarly organized whole genome table is not shown).

**Figure 3.**
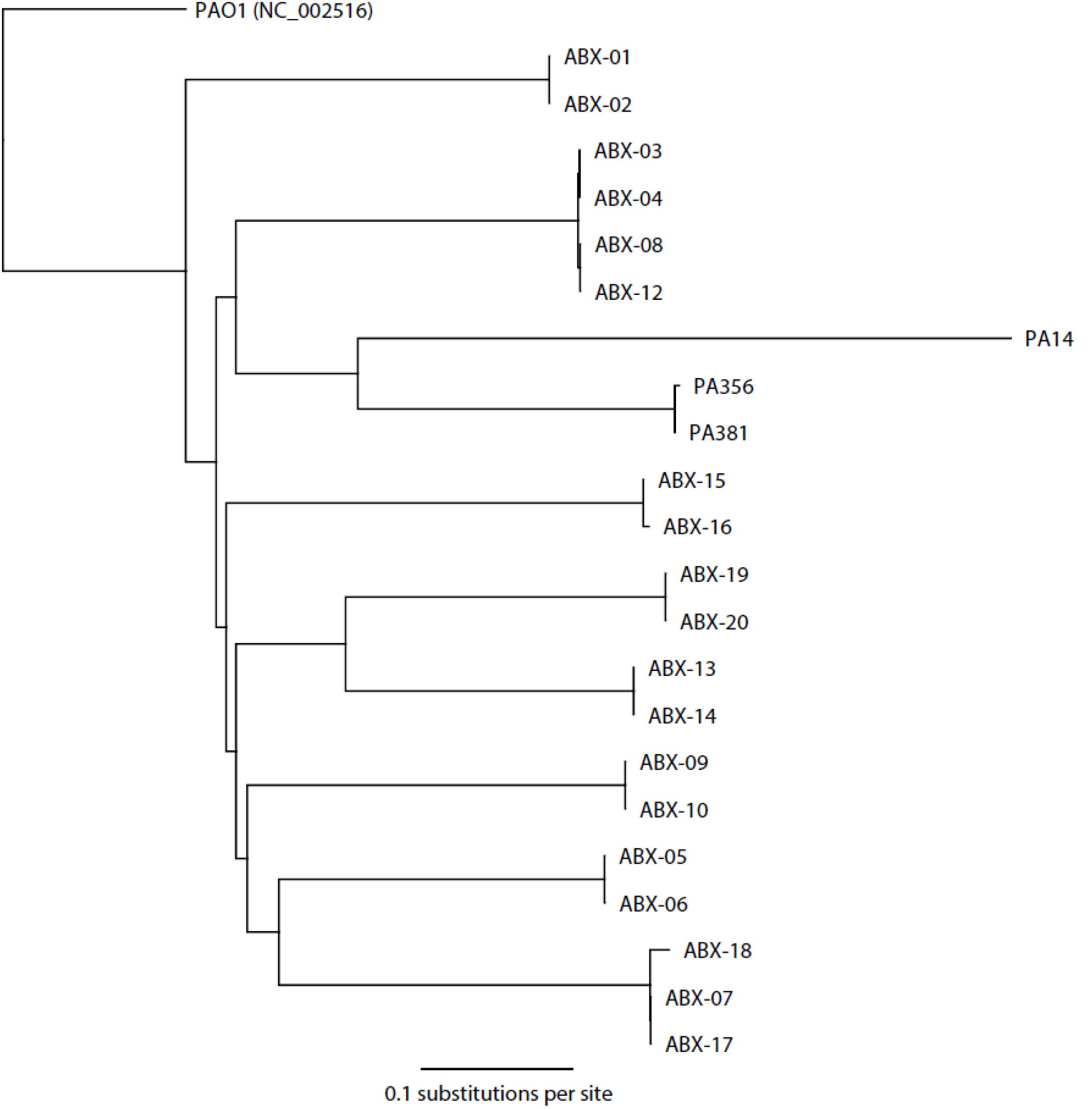
Whole genome tree Pan genome tree

The same antibiotic resistance genes well-known to PAO1 were found in each of the isogenic pairs of isolates sequenced in this study. A copy each of *ampC* and *bla_oxA-_*_50_*, aph(3’)-IIb, catB*7, and *fosA* were found to be conserved in all 19 isolates, identical to those of PAO1 (data not shown). A frameshift *“T”* insertion mutation at codon 293 of the 296-amino acid *ampR* was detected along with 3 other amino acid substitutions in both ABX-09 and −10, which were the only differences detected among all 19 isolates otherwise no concern for diverse MICs (Table 4). (Suppl Table 3). Based on genome coverage, only a 390kb region of ABX-01 showed a 3-fold increase in coverage relative to the background genome. This fragment aligned to a 350 kb region of PAO1 chromosome and 40 kb of the FluMu prophage present in *Pa* ATCC 27853 strain (CP015117.1). No other areas of significant coverage variation were observed in other strains.

The widely divergent MICs to CIP were most obvious in co-isolate pair ABX-01 and −02 and coisolate pair ABX-15 and −16 (Table 4). The classic mutations in the quinolone resistance-determining regions (QRDR) in *gyrA* codon 83 (T83I or T83A) were detected in 5 strains (ABX-01, −06, −08, −12, and −16), in which only two strains, ABX-01 and −16, produced an ultra high MIC of >32 μg/μl while others produced elevated MICs of 1-6 μg/μ1 (Table 4–5). A lesser known QRDR mutations in *gyrB* (S466F and S466Y) were also identified in strains ABX-18, −19, and −20, contributing to possibly elevated CIP MICs at 1,1, and 6 μg/μl respectively Table 2 and 4)(31). In fact, none of the amino acid substitutions (absent in PA14) in these four *(gyrA, gyrB, parC*, and *parE)* genes was able to entirely account for either the ultra-high or ultra-low MICs to CIP measured *in vitro* (Table 4–5).

Gene determinants associated with outer membrane porin proteins, efflux pumps, and their regulators were highly polymorphic amongst all isolates (Table S1 and S2). However, polymorphic mutations in this group of porin and efflux pumps hardly confirmed any previously known mechanisms of resistance. For example, despite their widely divergent MICs *in vitro*, the well known porin D determinant *oprD* (443 amino acid residues in PAO1) was similarly altered among 15 ABX isolates (ABX-3, −4, −5, −6, −8, −12, −13, −14, −15,-16, −7, −17, −18, −19 and −20), as well as reference strains ^*^356 and ^*^381; all isolates contained 21 identical non-synonymous substitutions along with a well-known loop L7 variant (Table S1) (30, 32). Above all, an OprD truncation was found in both ABX-19 and ABX-20 due to a stop codon at residue 176 without uniform measurements in their MIC levels against carbapenems or SMX-TMP (Table 4 and Suppl Table 1). For the rest 4 isolates belonging to two isogenic *Pa* pairs with MIC divergence, ABX-01 and −02 both contained three amino acid substitutions identical to those in PA14, while ABX-09 and −10 both contained a unique R310S (TableS1).

Isolates of ABX-01 and −02 both showed ultra high MIC of >32 μg/μl to SMX-TMP that could be correlated to possible *mexAB-oprM* overexpression resulting from frame shift deletions in their negative regulator gene *nalD* at codon 85 (-TGCGCTCGCTCTAC) and at 133 (-TG) respectively (33). However, the widely divergent MICs against all beta-lactam agents (AT, CX, PM, PTc, IP, and MP) found in ABX-01 and −02 could not be simply delineated by their presumed intact *oprD* but additional isogenic mutations in other relevant genes and regulators (Table 4 and Suppl Table 2). In fact, multiple mutations in efflux gene(s) (e.g. *mexAB-oprM, mexCD-oprJ, mexEF-oprN, mexXY-oprA, mexJK*, and multiple regulators *mexR, mexZ, nalC*, and *nalD*, etc.) were detected in these 19 isolates (Table 4–5, Table S2).

**Table 5.**
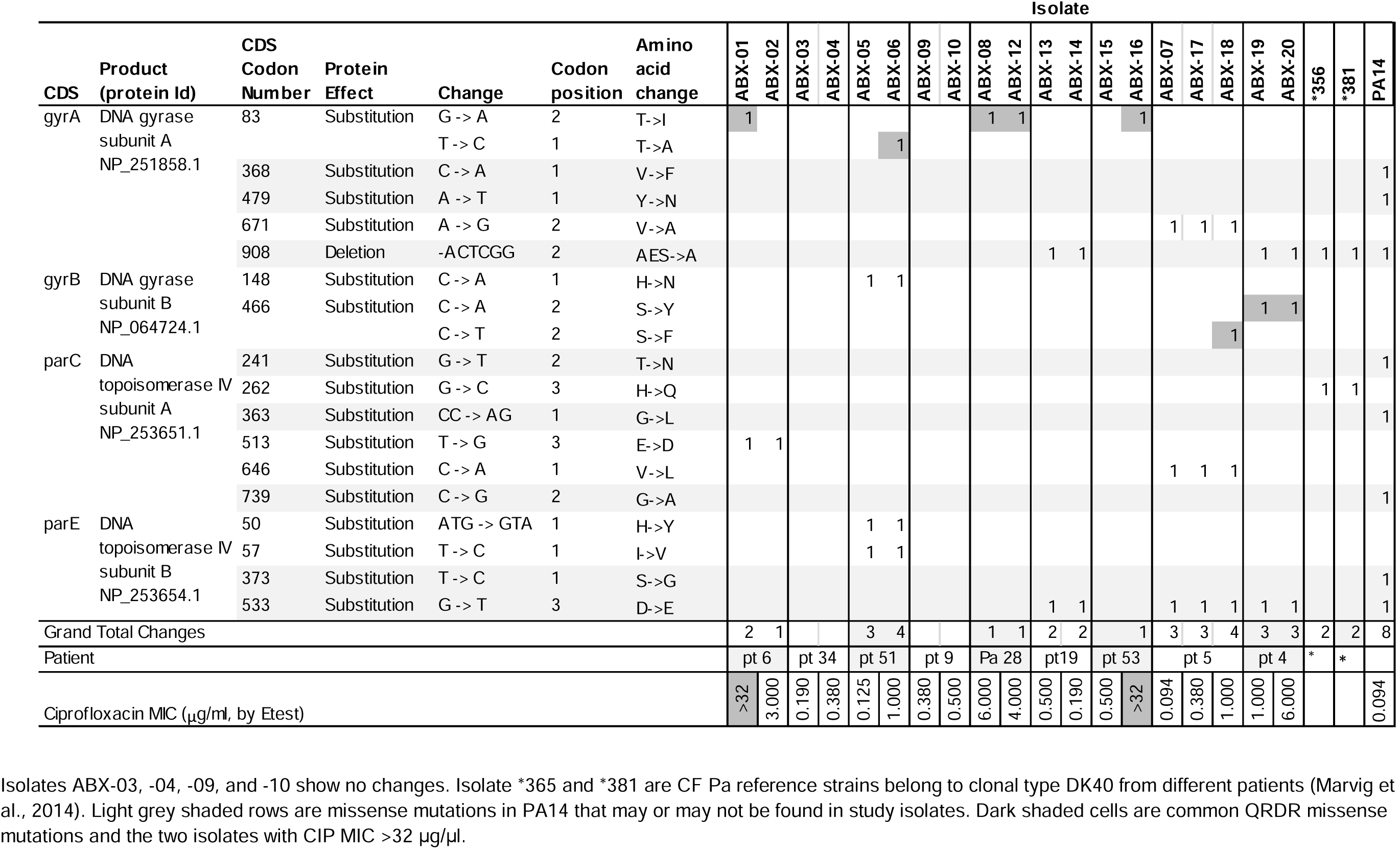
Polymorphic gene determinants associated with *P. aeruginosa* susceptibility to ciprofloxacin.

Mutations that result in non-synonymous substitutions were not only common in DNA mismatch repair genes (*mutL* and *mutS)*, but other determinants for nucleotide synthesis (*mutM* and *mutY)*, DNA repair or DNA/RNA folding, and recombination functions (*recB, recC, recD, recF, recG, redl, recN, recO, rdgC, rarA, recX, recR, uvrA, uvrB, uvrC*, and *uvrD)* in all ABX isolates and the CF *Pa* isolates ^*^356 and ^*^381 (Table S4). A 13-bp deletion in *mutS* (Table S4) was found only in ABX-16, but this isolate did not have an increased number of significant mutations in CDSs when compared to its isogenic ABX-15 or the rest of 17 ABX strains.

In order to get a rough estimate on functional CDSs that were immediately dispensable during *Pa* host specialization, we narrowed the analysis down to 627 mutually non-exclusive annotated CDSs with significant mutations, including CDS deletions, insertions, frameshift or nonsense/stop codon truncations among CF isolates, while excluding all hypothetical proteins, mutations that only resulted in amino acid nonsynonymous substitutions, and 319 CDS with polymorphisms that were in common to PA14 or present in PA14 only. These significant CDS mutations could potentially affect many aspects of bacterial cellular functions, such as metabolic functions, ABC transporter proteins, electron transport chain functions for ATP synthesis, iron acquisition, two-component sensor and regulators, the RNDs and membrane porin proteins, transcription regulators, type III secretion systems, DNA repair, RNA polymerization, structures of flagella and fimbriae, as well as biosynthesis of alginate, pyoverdine, sugar, lipids, nucleotides, amino acids and vitamins.

## Discussion

Our study demonstrated that antimicrobial MIC heterogeneity is unique to CF *Pa.* We found that the average number of *Pa* co-isolates per culture and average number of agents involved in discordant susceptibilities between co-isolates are both significantly higher in CF cohort than those of non-CF. Despite their significantly reduced susceptibilities to 5 of the 11 agents tested based on clinically established MIC breakpoints, isolates of CF *Pa* are likely to generate not only ultra-high but ultra-low MICs against a number of agents, (7, 34). Such heterogeneous and widely dispersed MIC patterns were associated with CF *Pa* affecting 9 out of 11 agents. Based on histograms and quantile-quantile plots, isolates of CF *Pa* show little resemblance to their non-CF *Pa* counterpart. With the genomic evidence of on average of 18% CDSs mutations in the 19 CF *Pa* genomes, we believe that the divergent MICs along with other structural and functional diversities are the net results of combined impact of these mutations in CF *Pa.* The distinct biological changes associated with CF *Pa* once again represent an evolutionary novelty due to their prolonged confinement to a specific host airway environment with selective pressure (3).

The current state of clinical laboratory susceptibility report has strictly followed the conventions of the three categories of interpretations for *in vitro* antimicrobial activities - susceptible, intermediate, and resistant. These interpretations have played and will continue to play significant roles in providing clinical reference values and guiding treatment decisions. CLSI has also developed practice guidelines using specific methods to provide “inferred mechanism of resistance” to call out specific types of resistance, such as “extended spectrum cephalosporin resistant *Enterobacteriaceae”* or staphylococcal “inducible clindamycin resistance” (21). These guidelines are highly timely and accurate for rapidly growing bacterial strains that are clonally homogeneous due to their brief encounters with the host organisms in settings of “acute infections”. However, the same laboratory standards have limitations to provide treatment references in areas of “chronic infections” where isolation of heterogeneous bacterial isogenic colony variants with discordant susceptibilities and/or production of heteroresistance patterns are common (Table 3–5, Fig. 1).

We prefer the term “heteroresistance” be used for two different situations: 1) co-isolates of the same bacterial species (often isogenic colony variants) producing discordant susceptibility patterns (Table 3–5), and 2) a seemingly “pure” and homogeneous isolate generating heterogeneous growth characteristics with reduced bactericidal effects in the presence of increasing antimicrobial drug concentrations *in vitro* (termed “Eagle-type” resistance, Fig 1) (19, 35). These heteroresistant patterns (Fig. 1 a-d) are most frequently found in bacterial isolates from CF airway specimens or from other localized chronic infections (19, 20). We therefore propose the use of Etest methods or broad-range micro dilutions for susceptibility testing of bacterial isolates exhibiting slow and heterogeneous growth *in vitro.* We also propose to introduce the term of “heteroresistance” in situations of an isolate producing non-homogenous susceptibility patterns where a MIC for the predominant growth is reported along with a descriptor (Fig 1A and B). Moreover, a term such as “avoid” may be more appropriate in situations of “eagle effect” such as those in Fig 1c-d. In situations of heteroresistance due to *Pa* co-isolates generating discordant results, laboratory interpretation and report could benefit from a specific “script” addressing the presence of isogenic isolates or clonal associates with divergent susceptibility patterns. Proper interpretive criteria and/or cross disciplinary accepted terminologies are critically important for the treatment of bacterial chronic infections.

Convergent mutational changes are not random among CF *Pa* genomes (e.g. *oprD, mexAB, mutL, mutS*, and electron transport complex E) from distantly related strains (Fig 3), which suggest their responses to a common selective pressure (23). Parallel to their common responses, the unshared changes detected between isogenic strains are indicative of pseudomonal transition to a multicellular lifestyle that gives rise to functional divergent but obligate syntrophic associates (3). The plethora of CDS polymorphisms (Table 4, Table S1-S4) detected in CF *Pa* are suggestive of not only their divergent and heteroresistance patterns but also a wide range of biological properties altered in these niche specialized bacteria (Fig 1) (3, 7).

The overall lake of genetic fluidity in ABX isolates is shown by no evidence of areas of sequences with increased coverage (except for ABX-01) for the presence of plasmids or other accessary genes that usually drives bacterial “superbugs” in free-living bacteria (36, 37). The 19 CF *Pa* representative strains all contain a chromosomal *ampC*, which upon induction is able to hydrolyze cephalosporins but not carbapenems while *blaOXA*-50 is a constitutively expressed oxacillinase (38). The spectrum of *aph(3′)-IIb* is known to confer resistance to a kanamycin and neomycin, but not AM, GM, or TM (39). In fact, none of the chromosomal resistance gene determinants (*ampC, blaOXA-*50, and *aph(3′)-IIb)* or the putative regulator AmpR mutations can be used to explain the phenomenon of widely divergent MICs generated from CF *Pa* isolates against any single agent or any single class of agents (Table 4–5, Table S3). Similar to the 47 *Pa* isolates reported by Lee et al that produced CIP MIC levels of 2-64 mg/L with only the *gyrA* T83I as the “first step” mutations (31), the 19 ABX isolates containing either the well defined QRDR mutations or novel amino acid substitutions did not show clear correlation with their wide range of CIP MICs consistently. Quinolone resistance in *Pa* has been well characterized to involve efflux mechanisms as well (40, 41). The highly polymorphic changes associated with the 19 ABX isolates in porin D, along with the RND family of efflux proteins and regulators showed no correlation to previously reported drug transporter or efflux pump activities affecting *Pa* susceptibilities uniformly to SMX-TMP, AT, carbapenems, or other classes of antimicrobials (26, 28, 29, 42, 43).

There is ample evidence of relaxed DNA mismatch repair systems due to mutations in both *mutL* and *mutS* in CF *Pa* isolates contributing to “hypermutability” and thus altered virulence and susceptibilities (7, 34, 44–46). CF *Pa* hypermutable state may not be limited to diverse mutations in *mutL* and/or *mutS* only, as a deleterious frameshift *mutS* mutation in ABX-16 did not show increased number of CDSs affected over other ABX isolates (Table 4 and Table S1-S4). Our additional findings of mutations in determinants responsible for nucleotide synthesis, DNA/RNA modification and recombination enzymes further support the long standing observation of the “genome decay” phenomenon as part of bacterial symbiotic response to host selective pressure (Table S4)(3, 23, 47). The scope and the scale of genome-wide mutations that specifically inactivated or altered the aspects of CF *Pa* structural and cellular functions may show us the way for the construction of a non-virulent pseudomonal symbiont that may be tailored to keep not only the wild-type *Pa* but other harmful organisms from colonizing (Table S1-S4)(3).

This study is limited by the lack of information regarding the duration of *Pa* infection in either cohort. Neither patient underlying diseases nor their treatment history were included in this analysis. The non-CF cohort of *Pa* isolates may have included patients whose pseudomonal infections are similar to patients with CF, or patients who have had prolonged infections affecting other organ or tissue systems. Future studies separating *Pa* isolates from newly infected host versus isolates from chronic infections (both CF and non-CF) would improve data clarity. Bacterial whole genome analysis from longitudinal isolates will be able to recapitulate a process with which a continuous host selected genome decay giving rise to a “host tissue-definitive” *Pa* specialist can be followed.

CF *Pa* and other specialists regularly escape from bacterial complexes to colonize a niche and, therefore, earn species name (3, 48). The inherently taxonomic based naming and reporting of a bacterial species without biological interpretation is no longer serving the reference role of *in vitro* testing or the medical need of the patients with CF or other chronic infections. With meaningful separation between homogeneous versus heterogeneous growth, or concordant versus discordant susceptibility profiles among isogenic co-isolates, the role of antibiotic treatment efficacy can then be studied and be used to compare outcomes. Clinical acknowledgement and understanding of laboratory findings of bacterial heteroresistance in the setting of chronic focal infections can open up opportunities to alternative therapeutic approaches (3).

## Materials and Methods

### Heteroresistant Isolates

To illustrate specific examples of heteroresistance at the strain level, we selected 4 representative CF *Pa* isolates that were identified at Seattle Children’s Hospital between 2011 and 2013. The antimicrobial MICs against Pa were determined and reported using our established consensus criteria that generally accept bactericidal agents such as beta-lactams and aminoglycosides at the Etest MIC cut point of no growth. For the bacteriostatic agent SMX-TMP, the Etest MIC at the point of 80% growth inhibition is accepted.

### Bacterial isolates from cultures included for antimicrobial MIC analysis

This analysis included all *Pa* isolates identified at Seattle Children’s Hospital (SCH) over a 7-month period (June 2013 to December 2013) and was approved by the SCH Institutional Review Board. All *Pa* isolates were identified by MALDI-TOF (Bruker Daltonics Inc.). Culture methods for isolates from CF patients were distinct from those used with non-CF isolates (7, 49). In order to minimize bias in data analysis for the purpose of this study, repeat isolates from the same patients in the study period were all included.

### Antibiotic susceptibility testing and MIC determination

Clinical samples were processed in the SCH Microbiology Laboratory according to CLSI guidelines. Antibiotic susceptibility was determined by E-test (bioMérieux) MIC method against 11 antimicrobial agents: amikacin, gentamicin, tobramycin, aztreonam, ceftazidime, cefepime, imipenem, meropenem, piperacillin/tazobactam, ciprofloxacin, and sulfamethoxazole-trimethoprim. Mueller Hinton (MH) agar plates (Remel, United States) were prepared by inoculating 0.5 McFarland saline suspensions of maximally purified and seemingly homogenous isolates before the application of Etest strips (Remel). The test plates were incubated at 35°C in ambient air normally for 18 to 24 hours for the determination of Etest MICs, but prolonged incubation (up to 36-48 hours) is common when testing CF isolates in order to achieve visual growth for MIC determination (7). The number of *Pa* co-isolates per culture was compared between CF versus non-CF cultures (Table 1). Antimicrobial susceptibility were interpreted using CLSI M100-S23 MIC breakpoint criteria (50). The percent susceptible *Pa* isolates was calculated as isolates that were interpreted as susceptible over the total number of *Pa* isolates. Discordant susceptibilities were defined as pseudomonal co-isolates that produced MIC differences meeting the interpretive criteria as susceptible and resistant to at least one of the 11 antibiotic agents in the same culture.

### Statistics

We assigned detection limits to those MIC data points for which the highest or the lowest detection limits were reached (e.g. an MIC value of >256 was entered as 256). We then applied logarithmic transformation to all the MIC readings (logMICs) to make the data less skewed for analysis. All subsequent analyses were based on logMICs.

We summarized number of *Pa* isolates per culture and average number of agents involved in discordant susceptibilities per culture using means and standard deviations, and compared them between CF and non-CF groups using t-tests. The percent susceptible value for each antimicrobial was compared between CF and non-CF isolates using Chi-squared test.

In the main statistical analysis we focused on the variance as the measure of dispersion. We first visually examined the distributions of the MIC readings between CF and non-CF isolates using histograms, density plots, and quantile-quantile plots. We then applied a robust test for equality of variances to examine the homogeneity of variances of MICs between isolates from CF and non-CF patients (51, 52). This test has demonstrated robustness even under nonnormality. For this analysis we treated data points as independent observations, even though some of the readings were from the same patients. All statistical analyses were performed using Stata (version 12.1; Stata Corp., College Station, TX).

### Bacterial whole genome sequencing

A subset of co-isolates was selected for whole genome sequencing based on their widely divergent *in vitro* antimicrobial MICs to at least one of the 11 agents tested (Table 4). The control genomes PA01 (GenBank Accession NC_002516.2), UCBPP-PA14 (PA14, GenBank Accession NC_008463.1) and two previously sequenced genomes of clonally related CF *Pa* reference strains ^*^356 and ^*^381 (GenBank SRA accession ERS402683 and ERS402710 respectively) were included for this analysis (23). PAO1 and PA14 are used to represent the two highly dissimilar *Pa* genomes for analysis as non-CF *Pa* references or “wild-type” susceptibility controls (7, 34, 53). All sequence polymorphisms associated with the ABX strains, strain ^*^356 and ^*^381, as well as PA14 were generated in reference to the PA0l genome (23). In this study, polymorphisms found in PA14 in reference to PAOl are assumed to have minimal functional impact in structural or biological functions including *in vitro* susceptibilities if both are treated as non-CF “wild-type” strains (Table 4, 5, suppl Table 1-4).

DNA was extracted from *Pa* isolates and diluted to 1ng/uL for sequencing library generation with third-volume Nextera XT and 14 cycles of PCR amplification (54, 55). Sequencing reads were adapter and quality trimmed using cutadapt, de novo assembled using SPAdes v3.9, and annotated using prokka (56, 57) (NCBI BioProject PRJNA359499). Reads were mapped to the PAO1 reference genome (NC_002516.2) and variant call files were generated using Geneious v9.1 using cutoffs of at least 7X coverage and 70% allele frequency. Sequences were deposited in NCBI Genbank under accessions SAMN06289359-SAMN06289367.

## Acknowledgements

We would like to thank Anne Marie Buccat and Jenny R. Stapp for their years of contributions to Cystic Fibrosis Microbiology Reference Laboratory operation and quality maintenance of clinical and research data. We would like to thank Seattle Children’s Microbiology team for their high quality performance and quality documentation of susceptibility testing and quality control results. Dr. Shuhua Yuan was a pediatrician from Shanghai Children’s Medical Center on Project Hope scholarship for her Microbiology and Infectious Diseases trainings at Seattle Children’s Hospital during August 2015 - July 2016.

## Author contributions

XQ, CZ, AG, and DM designed the research. CZ, AA, AG, and SY performed research and analyzed data. XQ, AA, AG, CZ and DM wrote the paper.

---

**Supplemental Table 1.** RND and Porin changes in P. aeruginosa isolates

**Supplemental Table 2.** Chromosomal amp*C* and *ampR* gene changes in *P. aeruginosa* isolates.

**Supplemental Table 3.** Polymorphic changes in genes governing DNA mismatch repair, nucleic acid synthesis, DNA or RNA modification and integrity, and DNA recombination functions in *P. aeruginosa* isolates.

**Supplemental Table 4.** Annotated CDSs with significant mutations, such as deletions, insertions, frame shift or nonsense/stop codon truncations among CF *P. aeruginosa* isolates (CDSs excluded-- all hypothetical proteins, mutations that resulted in amino acid nonsynonymous substitutions and CDS with polymorphisms that were in common to PA14 or present in PA14 only).

## References

1. Hentzer M, et al. (2003) Attenuation of Pseudomonas aeruginosa virulence by quorum sensing inhibitors. EMBO J 22(15):3803–3815.

2. Mena A, et al. (2007) Inactivation of the mismatch repair system in Pseudomonas aeruginosa attenuates virulence but favors persistence of oropharyngeal colonization in cystic fibrosis mice. Journal of bacteriology 189(9):3665–3668.

3. Qin X (2014) Chronic pulmonary pseudomonal infection in patients with cystic fibrosis: A model for early phase symbiotic evolution. Crit Rev Microbiol.

4. Oliver A, Baquero F, & Blazquez J (2002) The mismatch repair system (mutS, mutL and uvrD genes) in Pseudomonas aeruginosa: molecular characterization of naturally occurring mutants. Mol Microbiol 43(6):1641–1650.

5. Ciofu O, Riis B, Pressler T, Poulsen HE, & Hoiby N (2005) Occurrence of hypermutable Pseudomonas aeruginosa in cystic fibrosis patients is associated with the oxidative stress caused by chronic lung inflammation. Antimicrob Agents Chemother 49(6):2276–2282.

6. Oliver A, Canton R, Campo P, Baquero F, & Blazquez J (2000) High frequency of hypermutable Pseudomonas aeruginosa in cystic fibrosis lung infection. Science 288(5469):1251–1254.

7. Qin X, et al. (2012) Pseudomonas aeruginosa Syntrophy in Chronically Colonized Airways of Cystic Fibrosis Patients. Antimicrob Agents Chemother 56(11):5971–5981.

8. Proctor RA, et al. (2006) Small colony variants: a pathogenic form of bacteria that facilitates persistent and recurrent infections. Nat Rev Microbiol 4 (4) 295–305.

9. Wolter DJ, et al. (2013) Staphylococcus aureus small-colony variants are independently associated with worse lung disease in children with cystic fibrosis. Clinical infectious diseases: an official publication of the Infectious Diseases Society of America.

10. Grant SS & Hung DT (2013) Persistent bacterial infections, antibiotic tolerance, and the oxidative stress response. Virulence 4(4):273–283.

11. Anderson SW, Stapp JR, Burns JL, & Qin X (2007) Characterization of small-colony-variant Stenotrophomonas maltophilia isolated from the sputum specimens of five patients with cystic fibrosis. Journal of clinical microbiology 45(2):529–535.

12. Evans TJ (2015) Small colony variants of Pseudomonas aeruginosa in chronic bacterial infection of the lung in cystic fibrosis. Future Microbiol 10(2):231–239.

13. Sendi P, et al. (2010) Escherichia coli variants in periprosthetic joint infection: diagnostic challenges with sessile bacteria and sonication. Journal of clinical microbiology 48(5):1720–1725.

14. Grobner S, Beck J, Schaller M, Autenrieth IB, & Schulte B (2012) Characterization of an Enterococcus faecium small-colony variant isolated from blood culture. International journal of medical microbiology: IJMM 302(1):40–44.

15. Valderrey AD, et al. (2010) Chronic colonization by Pseudomonas aeruginosa of patients with obstructive lung diseases: cystic fibrosis, bronchiectasis, and chronic obstructive pulmonary disease. Diagn Microbiol Infect Dis 68(1):20–27.

16. Oliver A & Mena A (2010) Bacterial hypermutation in cystic fibrosis, not only for antibiotic resistance. Clin Microbiol Infect 16(7):798–808.

17. Pirnay JP, et al. (2009) Pseudomonas aeruginosa population structure revisited. PLoS One 4(11):e7740.

18. Pollack M ed (2000) Pseudomonas aeruginosa (Churchill Livingstone, New York, NY), 5th Ed, pp 2310–2327.

19. El-Halfawy OM & Valvano MA (2015) Antimicrobial heteroresistance: an emerging field in need of clarity. Clin Microbiol Rev 28(1):191–207.

20. Falagas ME, Makris GC, Dimopoulos G, & Matthaiou DK (2008) Heteroresistance: a concern of increasing clinical significance? Clin Microbiol Infect 14(2):101–104.

21. CLSI (Performance Standards for Antimicrobial Susceptibility Testing, in M100, ed Institute CaLS (Wayne, PA 19087).

22. CLSI (2009) Analysis and Presentation of Cumulative Antimicrobial Susceptibility Test Data. in M39-A3, ed Institute CLS (Wayne, PA 19087).

23. Marvig RL, Sommer LM, Molin S, & Johansen HK (2015) Convergent evolution and adaptation of Pseudomonas aeruginosa within patients with cystic fibrosis. Nat Genet 47(1):57–64.

24. Wilke MS, et al. (2008) The crystal structure of MexR from Pseudomonas aeruginosa in complex with its antirepressor ArmR. Proc Natl Acad Sci USA 105(39):14832–14837.

25. Daigle DM, et al. (2007) Protein modulator of multidrug efflux gene expression in Pseudomonas aeruginosa. Journal of bacteriology 189(15):5441–5451.

26. Kohler T, et al. (1996) Multidrug efflux in intrinsic resistance to trimethoprim and sulfamethoxazole in Pseudomonas aeruginosa. Antimicrob Agents Chemother 40(10):2288–2290.

27. Qin X, et al. (2008) Prevalence and mechanisms of broad-spectrum beta-lactam resistance in Enterobacteriaceae: a children’s hospital experience. Antimicrob Agents Chemother 52(11):3909–3914.

28. Braz VS, Furlan JP, Fernandes AF, & Stehling EG (2016) Mutations in NalC induce MexAB-OprM overexpression resulting in high level of aztreonam resistance in environmental isolates of Pseudomonas aeruginosa. FEMS Microbiol Lett 363(16).

29. Pan YP, Xu YH, Wang ZX, Fang YP, & Shen JL (2016) Overexpression of MexAB-OprM efflux pump in carbapenem-resistant Pseudomonas aeruginosa. Arch Microbiol 198(6):565–571.

30. Pirnay JP, et al. (2002) Analysis of the Pseudomonas aeruginosa oprD gene from clinical and environmental isolates. Environ Microbiol 4(12):872–882.

31. Lee JK, Lee YS, Park YK, & Kim BS (2005) Alterations in the GyrA and GyrB subunits of topoisomerase II and the ParC and ParE subunits of topoisomerase IV in ciprofloxacin-resistant clinical isolates of Pseudomonas aeruginosa. Int J Antimicrob Agents 25(4):290–295.

32. Epp SF, et al. (2001) C-terminal region of Pseudomonas aeruginosa outer membrane porin OprD modulates susceptibility to meropenem. Antimicrob Agents Chemother 45(6):1780–1787.

33. Evans K, et al. (1998) Influence of the MexAB-OprM multidrug efflux system on quorum sensing in Pseudomonas aeruginosa. Journal of bacteriology 180(20):5443–5447.

34. Wolter DJ, Black JA, Lister PD, & Hanson ND (2009) Multiple genotypic changes in hypersusceptible strains of Pseudomonas aeruginosa isolated from cystic fibrosis patients do not always correlate with the phenotype. J Antimicrob Chemother 64(2):294–300.

35. Eagle H & Musselman AD (1948) The rate of bactericidal action of penicillin in vitro as a function of its concentration, and its paradoxically reduced activity at high concentrations against certain organisms. J Exp Med 88(1):99–131.

36. Xiong J, et al. (2013) Complete sequence of pOZ176, a 500-kilobase IncP-2 plasmid encoding IMP-9-mediated carbapenem resistance, from outbreak isolate Pseudomonas aeruginosa 96. Antimicrob Agents Chemother 57(8):3775–3782.

37. Shen K, et al. (2006) Extensive genomic plasticity in Pseudomonas aeruginosa revealed by identification and distribution studies of novel genes among clinical isolates. Infection and immunity 74(9):5272–5283.

38. Girlich D, Naas T, & Nordmann P (2004) Biochemical characterization of the naturally occurring oxacillinase OXA-50 of Pseudomonas aeruginosa. Antimicrob Agents Chemother 48(6):2043–2048.

39. Zeng L & Jin S (2003) aph(3’)-IIb, a gene encoding an aminoglycoside-modifying enzyme, is under the positive control of surrogate regulator HpaA. Antimicrob Agents Chemother 47(12):3867–3876.

40. Llanes C, et al. (2011) Role of the MexEF-OprN efflux system in low-level resistance of Pseudomonas aeruginosa to ciprofloxacin. Antimicrob Agents Chemother 55(12):5676–5684.

41. Jacoby GA (2005) Mechanisms of resistance to quinolones. Clinical infectious diseases: an official publication of the Infectious Diseases Society of America 41 Suppl 2:S120–126.

42. Ocampo-Sosa AA, et al. (2012) Alterations of OprD in carbapenem-intermediate and -susceptible strains of Pseudomonas aeruginosa isolated from patients with bacteremia in a Spanish multicenter study. Antimicrob Agents Chemother 56(4):1703–1713.

43. Li XZ, Plesiat P, & Nikaido H (2015) The challenge of efflux-mediated antibiotic resistance in Gram-negative bacteria. Clin Microbiol Rev 28(2):337–418.

44. Lee DG, et al. (2006) Genomic analysis reveals that Pseudomonas aeruginosa virulence is combinatorial. Genome Biol 7(10):R90.

45. Lopez-Causape C, et al. (2013) Clonal dissemination, emergence of mutator lineages and antibiotic resistance evolution in Pseudomonas aeruginosa cystic fibrosis chronic lung infection. PLoS One 8(8):e71001.

46. Montanari S, et al. (2007) Biological cost of hypermutation in Pseudomonas aeruginosa strains from patients with cystic fibrosis. Microbiology 153(Pt 5):1445–1454.

47. Degnan PH, Yu Y, Sisneros N, Wing RA, & Moran NA (2009) Hamiltonella defensa, genome evolution of protective bacterial endosymbiont from pathogenic ancestors. Proc Natl Acad Sci U S A 106(22):9063–9068.

48. Georgiades K & Raoult D (2010) Defining pathogenic bacterial species in the genomic era. Front Microbiol 1:151.

49. Burns JL, et al. (1998) Microbiology of sputum from patients at cystic fibrosis centers in the United States. Clinical infectious diseases: an official publication of the Infectious Diseases Society of America 27(1):158–163.

50. CLSI (2009-2014) Performance Standards for Antimicrobial Susceptibility Testing. in M100-S19, ed Institute CaLS (Clinical and Laboratory Standards Institute, Wayne, PA 19087).

51. Brown MB& Forsythe AB (1974) Robust Tests for the Equality of Variances. Journal of the American Statistical Association 69(346):364–367.

52. Markowski CA & Markowski EP (1990) Conditions for the Effectiveness of a Preliminary Test of Variance. Am Stat 44(4):322–326.

53. Kos VN, et al. (2015) The resistome of Pseudomonas aeruginosa in relationship to phenotypic susceptibility. Antimicrob Agents Chemother 59(1):427–436.

54. Greninger AL, et al. (2015) Two Rapidly Growing Mycobacterial Species Isolated from a Brain Abscess: First Whole-Genome Sequences of Mycobacterium immunogenum and Mycobacterium llatzerense. Journal of clinical microbiology 53(7):2374–2377.

55. Herman EK, et al. (2013) The mitochondrial genome and a 60-kb nuclear DNA segment from Naegleria fowleri, the causative agent of primary amoebic meningoencephalitis. J Eukaryot Microbiol 60(2):179–191.

56. Bankevich A, et al. (2012) SPAdes: a new genome assembly algorithm and its applications to single-cell sequencing. J Comput Biol 19(5):455–477.

57. Seemann T (2014) Prokka: rapid prokaryotic genome annotation. Bioinformatics 30(14):2068–2069.

